# Temporal enhancement of cross-adaptation between density and size perception based on the theory of magnitude

**DOI:** 10.1101/2021.02.16.431522

**Authors:** Rumi Hisakata, Hirohiko Kaneko

## Abstract

The ability to estimate spatial extent is an important feature of the visual system. A previous study showed that perceived sizes shrank after adaptation to a dense texture and reported that this density–size aftereffect is modulated by the degree of density. In this study, we found that the aftereffect was also modulated by the temporal density of the adapting texture. The test stimuli were two circles, and the adapting stimulus had a dotted texture. The adapting texture refreshed every form 67ms to 500 ms, or not at all (static), during the adaptation. The results showed that the aftereffects from a refreshing stimulus were larger than those under the static condition. On the other hand, density adaptation lacked such enhancement. This result indicates that repetitive presentation of an adapting texture enhanced the density–size cross-aftereffect. The fact that density modulation occurs in both the spatial and temporal domains is consistent with the theory of magnitude, which assumes that the processing of the magnitude estimation of space, time, and numbers share a common cortical basis.

## Introduction

We can visually estimate distances between trees in a park even though there is nothing in that space. It is strange that our visual system can estimate “vacant space” despite it receiving no signal from that space. Concerning the estimation of vacancy, Hisakata, Nishida, and Johnston (2016) reported a new adaptation effect, called the density–size aftereffect, in which the perceived distance shrinks after adaptation to a dense texture [1]. They argued that this indicates that density plays a role as a metric for the estimation of distance between objects, and that adaptation to a dense texture reduces the scale, resulting in the distance between objects appearing smaller. In other words, apparent density mirrors the change in the scale of the metric itself. This property of adaptation is different from that of traditional size or spatial frequency adaptation, in which the direction of perceptual change is opposite to that of the adapting stimulus. Therefore, the representation of size or spatial frequency at early visual stages should remain unchanged even if the estimated distance between objects becomes shorter. In which case, at which stage is the visual metric read?

Concerning the estimation of space, time, and numerosity, Walsh (2003) proposed the theory of magnitude (TOM), a conceptual framework that assumes that the processing of magnitude estimation of space, time, and numbers share a common cortical basis [2]. Regarding the cortical area related to the TOM, the intraparietal sulcus (IPS) seems to be responsible for magnitude estimation because many physiological studies have shown that the judgement of size or orientation, such as the bisection task, and the estimation of lengths of line, numerosity, and duration all activate this region (e.g. [3–5]. Researchers have claimed that the magnitude estimation system in the IPS gathers sensory signals from cortical areas with different modalities and then makes a common representation of their magnitudes for action [2,6,7]. It is likely that this common magnitude processing system reads the special metric represented by density information in the perceptual system.

Concerning apparent numerosity, Burr and Ross (2008) reported that the apparent numerosity of texture decreases (increases) after adaptation to a large (small) number, indicating that the system for estimating numbers is adaptable [8]. Other studies showed that this aftereffect occurs in a higher cortical area, such as the IPS [3,9,10]; therefore, it is likely that a common system for magnitude estimation is adopted in the numerosity aftereffect. Recently, Aagten-Murphy and Burr (2016) reported that the numerosity aftereffect increases with repetition of the adapting stimulus, not with an increase in the adaptation duration [11]. This is noteworthy; the fact that the number of repetitions, not the duration, is crucial for the numerosity aftereffect indicates that apparent numerosity is represented in higher-order mechanisms.

In this study, we investigated the effects of both the repetition and duration of an adapting stimulus on the density–size aftereffect. If the density–size aftereffect demonstrated in Hisakata et al. (2016) shares common mechanisms with the numerosity aftereffect mentioned above, it should have a temporal property similar to that of the latter. In other words, the repetition, not the duration, of an adapting texture should increase adaptation-induced space compression, as with numerosity adaptation. In experiment 1, we measured the timecourse of the density–size aftereffect in a similar way as that employed by Aagten-Murphy and Burr (2016) for the numerosity aftereffect. We manipulated both the presented repetition number and duration of the adapting texture and measured the perceived size of an object before and after adaptation. In experiment 2, we directly examined the effect of repetition of the adapting texture on both size and density perceptions.

## Results

In experiment 1, we measured the perceived size of a circle before and after adaptation to examine the timecourse of the density aftereffect. During a session, two circles were presented in the left and right visual fields and an observer judged which circle was larger (Fig. 1a). In the first 60 trials, the test stimulus alone was presented repeatedly in the left and right visual fields for 200 ms (Fig. 1b). In the middle 60 trials, an adapting texture was inserted between successive test stimulus presentations. In the last 120 trials, again only the test stimulus was presented. One session was composed of one staircase, so that we could track the shift in perceived size during the session. We manipulated the duration and repetition number of the adapting stimulus to examine which factor was crucial for the aftereffect. We had two duration conditions, 1 or 5 s, and three repetition conditions, in which both the position and luminance of the texture were refreshed every 100 or 300 ms or were not refreshed (static).

**Figure 1.**
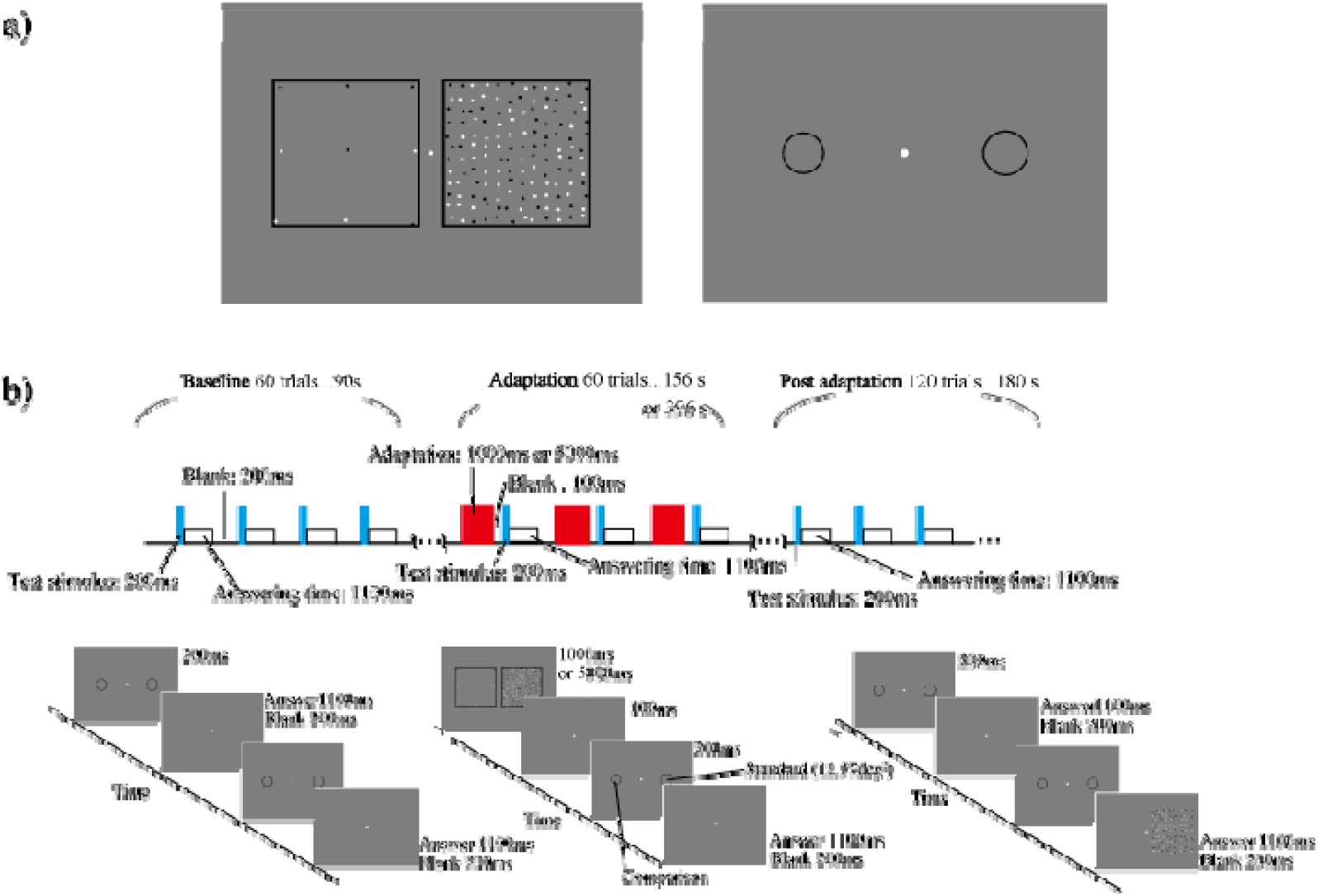
a) Example of an adapting texture (left panel) and the test stimulus (right panel). b) Sequence for one experimental session. In the baseline and post-adaptation trials, only the test stimulus was presented. In the adaptation trials, an adapting texture was presented between presentations of the test stimulus. The size of the comparison stimulus was manipulated according to the staircase method in a one-up, one-down manner.

Fig. 2a shows the changes in perceived size before, during, and after texture adaptation under all refresh conditions for each observer. Red and green lines in each panel show the results of the adaptation duration conditions of 1 and 5 s, respectively. The perceived size of a circle presented on the adapted side shrank in the adaptation phase under all conditions. Interestingly, the shrinkage reached the maximum value right after the beginning of the adaptation phase.

**Figure 2.**
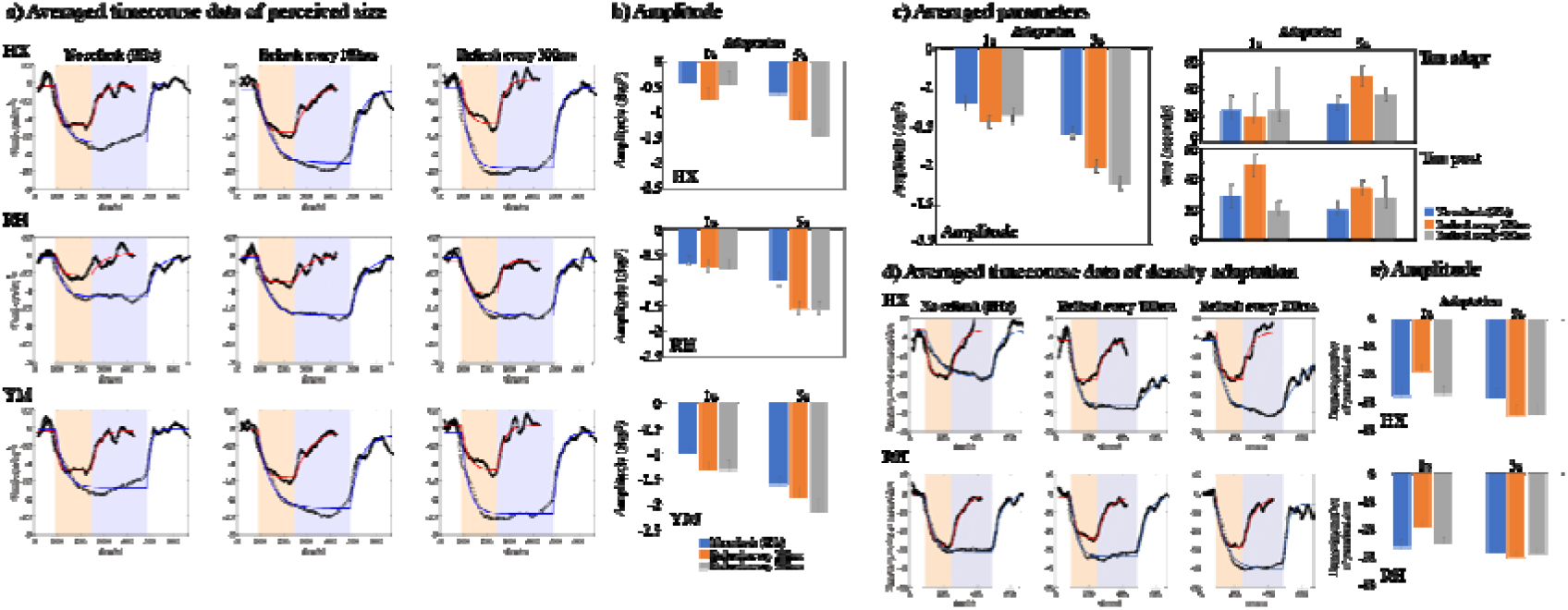
a) Averaged timecourse data of perceived size before, during, and after density adaptation for each observer (each indicated by a pair of letters). In (a) and (d), each panel shows the results under each refresh condition; light red and blue areas indicate the adaptation duration conditions of 1 and 5 s, respectively. Red and blue solid lines indicate functions fitted to the data of the adapting conditions, 1 and 5 s, respectively. b) Estimated amplitude of the density–size aftereffect with a bootstrap simulation with 10,000 runs. In (b) and (e), the amplitude was estimated using the median of the histogram from non-parametric bootstrapping. The error bars in (b) and (c) indicate 95% confidence intervals estimated with the percentile bootstrap method. c) Averaged parameters across the observers with the bootstrap simulation run 10,000 times. d) Averaged timecourse data of perceived density before, during, and after adaptation for each observer. e) Estimated amplitude of density–density aftereffects with a bootstrap simulation with 10,000 runs.

To show the timecourse of the adaptation effect quantitatively, we fitted a function (see Materials and Methods) and calculated the amplitudes and rise and decay times. After this fitting, we assessed whether differences were significant based on the overlap between 95% confidence intervals estimated with the percentile bootstrap method. The amplitudes of shrinkage were larger for the 5-s adaptation than for the 1-s one under all refresh rate conditions, showing that the adaptation duration is important for the density–size aftereffect. This result is inconsistent with the temporal characteristics of the numerosity aftereffect shown in Aagten-Murphy and Burr (2016) [11]. The amplitude of the aftereffect under the static (no refresh) condition was smaller than those with a refresh rate of once every 100 and 300 ms for both adaptation durations of 1 and 5 s (Fig. 2b). There results indicate that the refresh frequency of the adapting texture also affects shrinkage, especially under the 5-s adaptation condition.

To compare this characteristic of size perception with that of density perception, we conducted the same measurement using a textured pattern instead of a circle as the test stimulus. Here, the pattern consisted of dots in random positions, and the observer judged whether the left or right texture was denser. For density perception, the magnitude of the aftereffect did not differ between adaptation conditions of 1 and 5 s; furthermore, it did not seem that refreshing the adapting texture affected the magnitude of the aftereffect (Fig. 2d and e). This result suggests that the density–density aftereffect saturates even when the adaptation period is short (1 s), whereas the density–size aftereffect was affected by both the adaptation duration and refresh frequency, indicating that the magnitude of the density–size aftereffect is influenced by temporal adaptation.

To examine the effect of repetition on both the density–size and density–density aftereffects directly, we manipulated the repetition number in experiment 2. During this experiment, the first adaptation period lasted for 60 s, and top-up adaptation was presented for 5 s in each trial (Fig. 3a). Four staircases were run in each session, and the perceived size and density was defined by the average of the last 10 reversals of each staircase. The refresh rate of the adapting texture varied from 0 (static) to 15 Hz. Fig. 3b shows the changes in perceived size (left) and density (right) due to the aftereffect. Temporal refreshing of the adapting texture enhanced the aftereffect on size perception (*F*(4,28) = 10.07, *p* < 0.0001, *η*^2^ = 0.206, and Ryan’s method multiple comparison showed 0 Hz < other temporal frequencies); on the other hand, it did not change the aftereffect on density perception except under a refresh frequency of 15 Hz (*F*(4,28) = 41.29, *p* = 0.034, *η*^2^ = 0.099, Ryan’s method multiple comparison showed 0 Hz < 15 Hz). As in experiment 1, we found a difference in the characteristics of temporal enhancement for size and density perceptions after adaptation to a dense texture.

**Figure 3.**
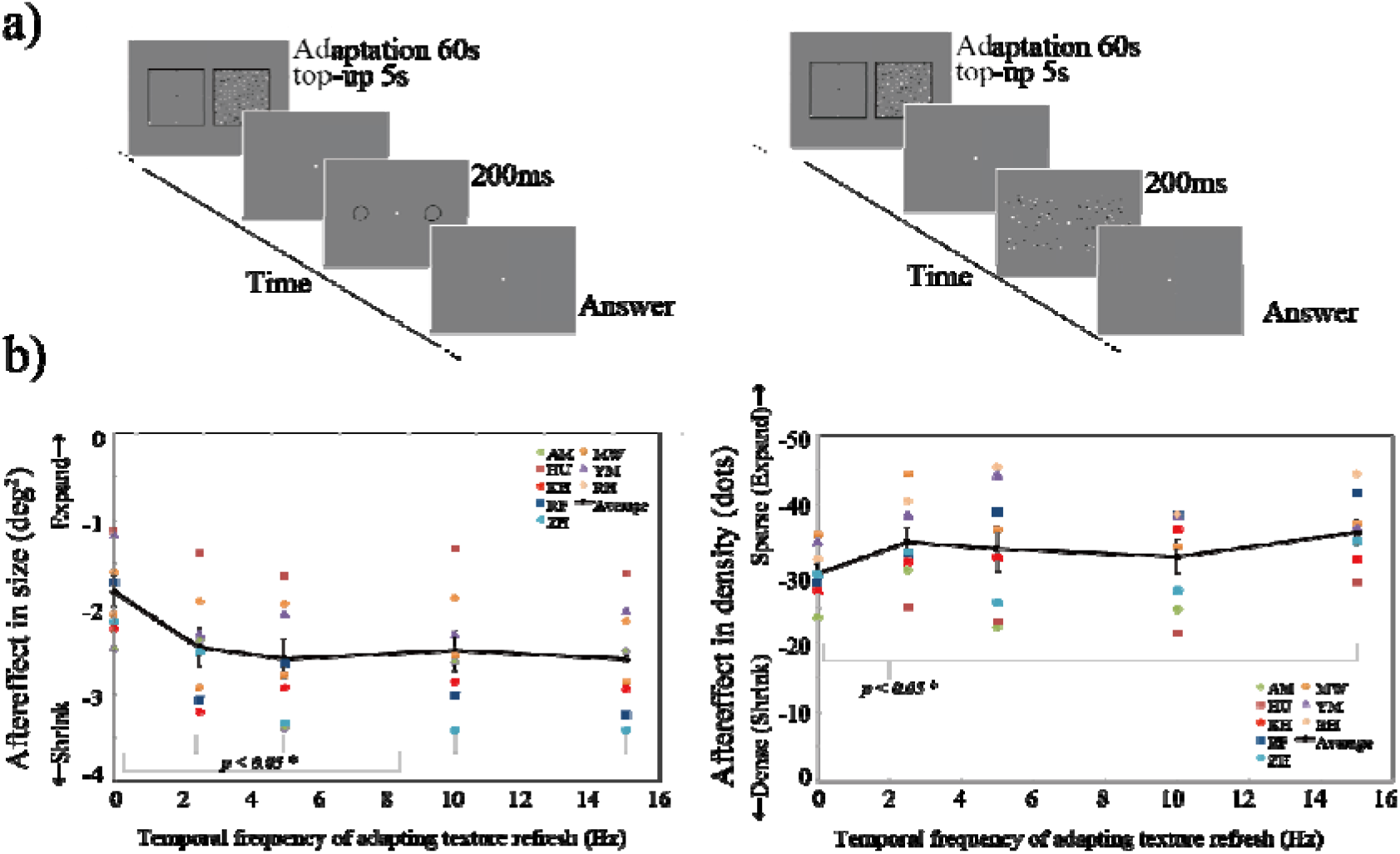
a) Stimulus presentation in experiment 2. b) Results of the aftereffect for each observer (each indicated by a pair of letters) including the average values. The left and right panels show the change in perceived size and density after adaptation, respectively, as a function of the refresh frequency of the adapting stimulus. Error bars indicate the standard errors of the means.

## Discussion

Based on the TOM, we hypothesized that the density–size aftereffect should be affected by temporal modulation of the adapting texture, as with the numerosity aftereffect. Here, we showed that repeated presentation of the adapting texture increased the shrinkage in perceived size after adaptation. We consider this an indirect effect of temporal adaptation at the higher stage of estimating a common “magnitude.” If all magnitudes of different perceptual modalities, such as time, space, and number, are estimated in a common cortical area, it is likely that the temporal characteristics of an adapting stimulus are similar for those modalities.

We assume that two stages are necessary to explain the density–size aftereffect. The first stage is density representation as a spatial metric in the visual sensory area. The second stage is the estimation processing based on that metric. The conceptual relationship between these stages is depicted in Fig. 4. As mentioned in our Introduction, the metric could change due to a long exposure to a dense texture; in other words, adaptation occurs at the visual processing stage. The estimation stage is then related to the representation of common magnitude, indicating that magnitude estimation in one modality could affect that in other modalities.

**Figure 4.**
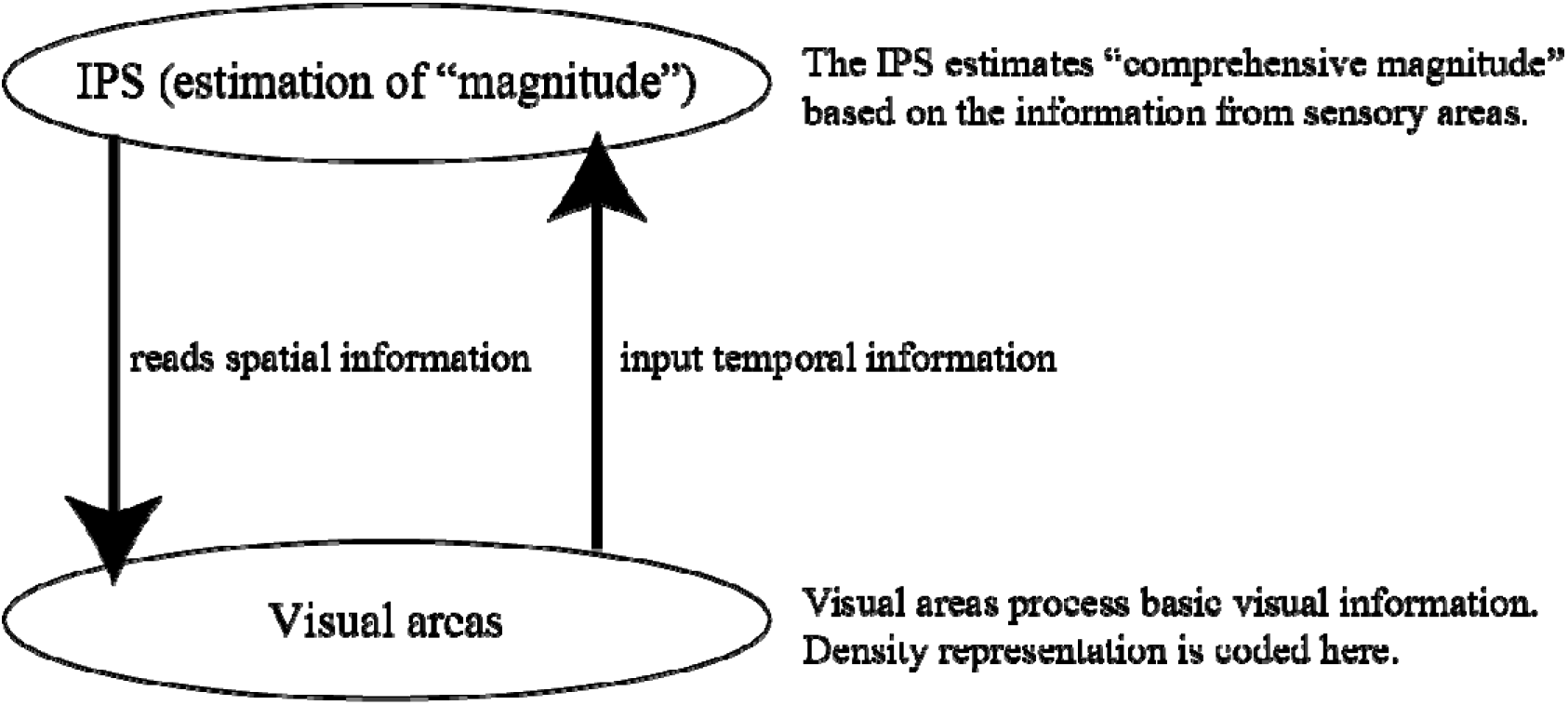
Conceptual model of the functional relationship between the intraparietal sulcus (IPS) and visual sensory areas.

A possible mechanism would be that a common system to code magnitude adapts to the temporal number of a particular presentation because it also estimates the temporal presentation number, and temporal duration could be coded there [12–14]. The temporal frequency and velocity of stimuli affect the perceived duration (e.g. [15–17]. For example, after a long exposure to flickering or moving stimuli with high temporal frequency, the magnitude estimation system is exhausted, and the output value would be smaller. If a common magnitude estimation system is used for spatial estimation, the perceived size or distance would be affected by temporal refreshing of an adapting texture, resulting in a density–size aftereffect, as with the perceived duration. We observed this effect in the present results. We suppose that this indirect effect of temporal refreshing on spatial estimation is limited; therefore, the effect was saturated at a refresh rate of 2 Hz. On the other hand, the estimation of density itself is represented at the earliest stage of visual processing, in which spatial and temporal information are processed independently. Therefore, temporal refreshing did not modulate the aftereffect on density perception. In experiment 2, refreshing at a relatively high frequency (15 Hz) induced a larger density–density aftereffect, whereas a low frequency did not. We speculate that the apparently high-density texture produced by refreshing with high frequency caused the greater density aftereffect. Because we used white and black dots on a gray background and the positions changed with a 15 Hz refresh rate, an afterimage might easily have remained (e.g. [18] and be treated as a member of the adapting dots, resulting in an increased density of the adapting texture.

In this study, we examined the relationship between density adaptation and the magnitude estimation system. We assumed that if the magnitude estimates of size and distance are related to the density–size aftereffect, temporal modulation of an adaptation that can affect other modalities, such as numerosity adaptation, should also affect the density–size aftereffect. Here, the results showed that the perceived size shrinkage after adaptation to a dense texture increased as the frequency of temporal refreshing of an adapting texture increased, whereas this was not the case for the density–density aftereffect. From these results, we propose that basic density representation in the visual area acts as a spatial metric and the magnitude estimation system reads this spatial metric. For our next step, using functional magnetic resonance imaging, we will identify the brain area related to magnitude and density representations after adaptation.

## Materials and Methods

### Experiment 1

Stimuli were generated on a computer (Apple MacBookPro 2016) and displayed on a liquid crystal display gaming monitor (EIZO FROIS 27-in, 2560 × 1440 pixels, refresh rate 100 Hz, 2.24 arcmin/pix, mean luminance 54.5 cd/m^2^, gamma-corrected). The viewing distance was 57.3 cm, and the size of the adapting texture was 15 × 15°. Three observers consented to participate in the experiment (one of the authors and two observers naïve to the purpose of the study). They all had normal or corrected-to-normal visual acuity. The study was approved by the Tokyo Institute of Technology Ethics Committee, Japan, and conformed to guidelines of the Declaration of Helsinki on the of human observers in research.

The adapting texture was composed of white and black dots with a diameter each of 10 pixels. The test stimulus was an open circle with a black edge. We used the simple staircase method (one-up, one-down) to measure the perceived size of the test circle. experimental session had one staircase for measuring the timecourse of the transition in perceived size. The session consisted of three parts: baseline, adaptation, and post-adaptation. In the baseline part, two circles were presented for 200 ms in both the left and right visual fields; one was the standard stimulus (12.25 deg^2^) and the other was the comparison stimulus (the size varied according to the observer’s previous response). Within 1100 ms after presentation, the observer responded with which circle, left or was larger. After a 200-ms blank, the next stimulus was presented. Each presentation of test stimuli (one trial) was repeated 60 times, taking a total of 90 s. During the periods, adapting textures were presented between presentations of the test circles. Textures were presented in both the left and right visual fields with black frames to prevent a size aftereffect. One of the textures consisted of 144 dots and the other Each dot in the adapting textures was positioned at a point on a square grid with a displacement of up to 30 arcmin, which was updated every 300 ms (3.33 Hz) or 100 ms (10 Hz), or not updated (static). There were two adaptation duration conditions (1 and 5 s). The adaptation part consisted of 60 trials and took 156 or 396 s depending on the stimulus duration. In the post-adaptation part, only the test stimulus was presented, as in the baseline part.

In each trial, we recorded the size of the stimulus compared to the standard one (12.25 deg^2^). The values of the first 10 trials were excluded from analysis because these included initial fluctuations. We defined the average value from the 11^th^ to 60^th^ trials as the baseline, and this baseline value was subtracted from all analyzed data. After subtraction, we calculated moving averages with a range of ±22 trials. We fitted the following function to the timecourse data with a non-linear least squares method:

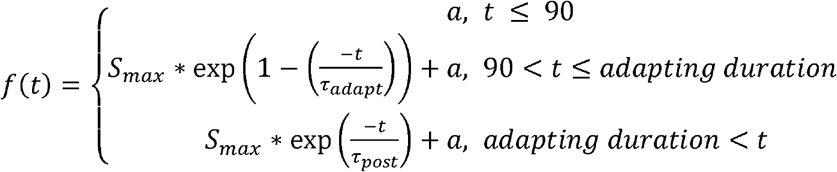

where *t* is the time in seconds, *a* is the baseline of shrinkage before adaptation, which should be approximately 0, *S*_*max*_ is the maximum value of the magnitude of the effect, *τ*_adapt_ is the time in which the maximum value is reached, and *τ*_post_ is the decay time. We defined the difference between *a* and *S*_*max*_ as the amplitude, which is plotted in Fig. 2. Non-parametric bootstrapping was carried out with 10,000 runs to calculate the 95% confidence intervals of these parameters with the percentile method.

We used the same procedure to measure density adaptation. In this case, only the test stimulus was different from the size trials, in that the texture consisted of white and black dots whose positions were randomly assigned within an area of 15 × 15°. The standard texture contained 49 dots, and the numbers of dots for estimating density varied according to the staircase method. The presentation duration of the test stimulus was 200 ms.

### Experiment 2

The apparatus and stimuli were the same as those used in experiment 1. Eight observers consented to participate in the experiment (one of the authors and seven observers naïve to the purpose of the study). The duration of the first adaptation period was 60 s with a 5-s top-up adaptation in subsequent trials. Each dot in the adapting texture was positioned at a point on a square grid with a random displacement of up to 30 arcmin, which was updated every 66.67 ms (15 Hz) or 300 ms (3 Hz), or not updated (static). The center of the stimulus was located at 9.55° in terms of visual angle from the fixation point. We used the one-up, one-down staircase method to measure the point of subjective equality. The position of the adapting visual field, left or right, was balanced among sessions. There were breaks of at least 5 min between sessions. We ran four sessions for each condition and defined the magnitude of perceived shrinkage using the average of the last 10 trials. For the density–density adaptation measurement, the procedure was the same except that the test stimulus was a texture composed of white and black dots each with a diameter of 10 pixels that were positioned randomly within the adaptation area.

## Author Contributions

Rumi Hisakata: Conceptualization, Methodology, Software, Investigation, Visualization, Writing-Original draft preparation, Writing-Reviewing and Editing, Funding acquisition. Hirohiko Kaneko: Conceptualization, Methodology, Visualization, Supervision, Writing-Reviewing and Editing.

